# Dissection of the ATPase active site of McdA reveals the sequential steps essential for carboxysome distribution

**DOI:** 10.1101/2021.04.02.438200

**Authors:** Pusparanee Hakim, Anthony G. Vecchiarelli

## Abstract

Carboxysomes, the most prevalent and well-studied anabolic bacterial microcompartment, play a central role in efficient carbon fixation by cyanobacteria and proteobacteria. In previous studies, we identified the two-component system called McdAB that spatially distributes carboxysomes across the bacterial nucleoid. McdA, a ParA-like ATPase, forms a dynamic oscillating gradient on the nucleoid in response to carboxysome-localized McdB. As McdB stimulates McdA ATPase activity, McdA is removed from the nucleoid in the vicinity of carboxysomes, propelling these proteinaceous cargos toward regions of highest McdA concentration via a Brownian-ratchet mechanism. However, how the ATPase cycle of McdA governs its *in vivo* dynamics and carboxysome positioning remains unresolved. Here, by strategically introducing amino acid substitutions in the ATP-binding region of McdA, we sequentially trap McdA at specific steps in its ATP cycle. We map out critical events in the ATPase cycle of McdA that allows the protein to bind ATP, dimerize, change its conformation into a DNA-binding state, interact with McdB-bound carboxysomes, hydrolyze ATP and release from the nucleoid. We also find that McdA is a member of a previously unstudied subset of ParA family ATPases, harboring unique interactions with ATP and the nucleoid for trafficking their cognate intracellular cargos.

## INTRODUCTION

The ParA family of ATPases play major roles in the subcellular organization of bacterial cells, with members involved in the positioning of a wide array of intracellular cargos including plasmids, chromosomes, the divisome, flagella, chemotaxis clusters, and carbon-fixing organelles called carboxysomes (Lutkenhaus, 2012; Vecchiarelli *et al.*, 2012; Kiekebusch and Thanbichler, 2014). How ATP is used to organize such a diversity of genetic and proteinaceous cargos remains unclear. ParA family members are defined by the presence of a deviant Walker A motif, along with Walker A’ and Walker B motifs (Koonin, 1993). Aside from these motifs that make up the ATP-binding pocket, few similarities exist at the sequence level. But structurally, all ParA family members solved to date form very similar nucleotide-sandwich dimers (Schumacher *et al.*, 2012, 2019; Zhang and Schumacher, 2017). ATP binding stabilizes dimerization because of an invariant “signature” lysine residue that defines the deviant Walker A box, which makes cross-contacts with the γ-phosphate of the opposing monomer making up the sandwich dimer (Dunham *et al.*, 2009).

The ParA family is named after its best-studied member. The ParA ATPase is part of tripartite DNA segregation system that <u>par</u>titions and positions replicated copies of chromosomes and low-copy plasmids to opposite cell halves, thus ensuring faithful inheritance of these genetic cargos after cell division (Baxter and Funnell, 2014; Badrinarayanan *et al.*, 2015; Jalal and Le, 2020). Cytoplasmic ParA monomers bind ATP and form the ATP-sandwich dimer (Davey and Funnell, 1997; Zhang and Schumacher, 2017). The ParA dimer then undergoes an ATP-specific conformational change that licenses binding to nonspecific DNA *in vitro*, which equates to binding the bacterial nucleoid *in vivo* (Hester and Lutkenhaus, 2007; Castaing *et al.*, 2008; Vecchiarelli *et al.*, 2010). In its DNA-binding form, ParA can robustly interact with its partner protein, ParB (Pratto *et al.*, 2008). ParB dimers site-specifically load onto the plasmid, or chromosome, to be partitioned via specific binding to a centromere-like site, typically called *parS* (Baxter and Funnell, 2014; Jalal and Le, 2020). ParB dimers spread from *parS* onto flanking DNA to form a massive multimeric nucleoprotein complex (Sanchez *et al.*, 2015; Funnell, 2016). This ParB-*parS* complex can interact with ParA dimers and stimulate its ATPase activity, which is coupled to ParA release from the nucleoid (Hwang *et al.*, 2013; Vecchiarelli *et al.*, 2013; Volante and Alonso, 2015). The resulting ParA depletion zone that forms around the ParB-*parS* complex also provides a ParA concentration gradient on the nucleoid. In this Brownian-ratchet mechanism, ParB-*parS* complexes on newly replicated chromosomes or plasmids are bidirectionally segregated to opposing cell-halves as they chase higher concentrations of ParA along the nucleoid in opposing directions (Vecchiarelli *et al.*, 2010, 2014).

A growing list of protein-based cargos have been shown to also require a ParA-type ATPase for their subcellular organization, including carboxysomes (Lutkenhaus, 2012; Vecchiarelli *et al.*, 2012). Carboxysomes are carbon-fixing organelles found in all cyanobacteria and most carbon-fixing proteobacteria (Turmo *et al.*, 2017), and are responsible for roughly a third of global carbon fixation (Cohen and Gurevitz, 2006). By encapsulating the enzymes Ribulose-1,5-bisphosphate carboxylase/oxygenase (Rubisco) and carbonic anhydrase in a selectively permeable protein shell, the resulting CO_2_-rich microenvironment within carboxysomes ensures that carboxylation of ribulose-1,5-bisphosphate is favored over the undesired process of photorespiration where O_2_ is fixed instead of CO_2_ (Kerfeld *et al.*, 2018). Despite the importance of carboxysomes to the global carbon cycle, the mechanisms underlying their subcellular organization remains unclear.

In 2010, Savage and colleagues showed that a ParA-like ATPase, now termed <u>M</u>aintenance of <u>c</u>arboxysome <u>d</u>istribution protein <u>A</u> (McdA), was required for the equidistant positioning of carboxysomes down the length of the rod-shaped cyanobacterium *Synechococcus elongatus* PCC7942 (henceforth *S. elongatus*) (Savage *et al.*, 2010). More recently, we found that McdA functions with a partner protein, called McdB, which associates with the carboxysome cargo and is required for the dynamic oscillatory behavior of McdA *in vivo* (MacCready *et al.*, 2018). ATP-bound McdA has non-specific DNA binding activity and McdB stimulates McdA ATPase activity as well as its release from a non-specific DNA substrate *in vitro*. From these biochemical findings, we proposed that McdB-bound carboxysomes locally stimulate the release of McdA from the nucleoid, and the resulting McdA gradients are then used to drive the movement and equidistant positioning of carboxysomes across the nucleoid region of the cell; akin to DNA partitioning by ParABS systems. However, it remains to be determined how the ATP cycle of McdA governs the molecular events required for its dynamic oscillatory patterning and the positioning of McdB-bound carboxysomes across the nucleoid.

There are notable differences that set *S. elongatus* McdA apart from classical ParA family ATPases. For example, the signature lysine residue that defines the ParA family is absent in the deviant Walker A box of McdA. Also intriguing was the finding that McdA possesses a substantially higher ATPase activity compared to ParA ATPases involved in DNA partitioning (Ah-Seng *et al.*, 2009; Vecchiarelli *et al.*, 2010; MacCready *et al.*, 2018). These differences drove us to dissect the molecular events of carboxysome positioning by McdA and identify how these steps are coupled to its ATP cycle.

Despite these differences, it was recently shown that an McdA homolog shares the adenine-nucleotide sandwich dimer structure solved for several other ParA family ATPases (Schumacher *et al.*, 2019) **(Figure 1A)**. Additionally, many of the invariant amino acids critical for ATP-dependent functions are also found in McdA; with the exception of the signature lysine residue in the Walker A box **(Figure 1B)**. To dissect how ATP-binding and hydrolysis mediates McdA function in carboxysome positioning, we introduced strategic amino acid substitutions in the ATP-binding pocket of McdA. The mutations are synonymous with “trap” mutants made in several well-studied ParA family members involved in the positioning of plasmids (Fung *et al.*, 2001; Libante *et al.*, 2001; Barillà *et al.*, 2005; Vecchiarelli *et al.*, 2010, 2013), chromosomes (Leonard *et al.*, 2005), the divisome (Lutkenhaus and Sundaramoorthy, 2003; Kiekebusch *et al.*, 2012; Schumacher *et al.*, 2017), flagella (Ono *et al.*, 2015; Schuhmacher *et al.*, 2015), and chemotaxis clusters (Roberts *et al.*, 2012; Ringgaard *et al.*, 2014) **(Summary in Figure 1C, detailed in Table S1)**. The data presented in this study connects key steps in the ATP cycle of McdA to the stepwise events required for distributing McdB-bound carboxysomes across the cyanobacterial nucleoid.

**Figure 1:**
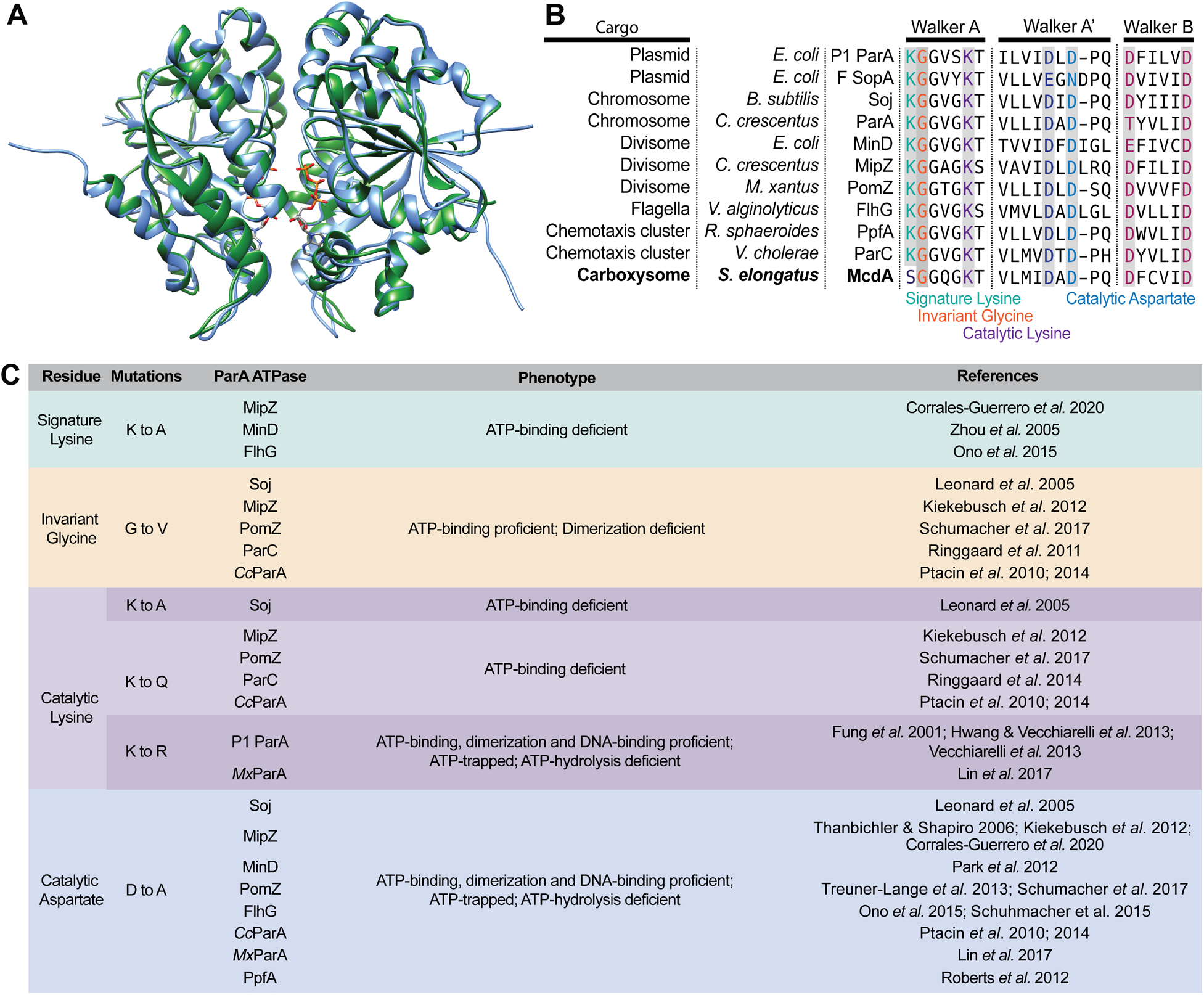
McdA shares structure and sequence conservation with ParA-type ATPases. (**A**) The crystal structure of *Cyanothece* McdA[D38A] (green; PDB entry 6nop) was superimposed on to the modelled structure of *S. elongatus* McdA (blue) with ATP molecules (sticks) in the sandwich dimer interface. (**B**) Amino acid sequence alignment of the Walker A, A’ and B motifs conserved among ParA family ATPases. Invariant residues are shaded grey. The signature lysine (green), invariant glycine (orange) and catalytic lysine (purple) in the Walker A motif and the catalytic aspartate in the Walker A’ motif were mutated in this study. (**C**) Summary of strategic mutations studied in ParA family members and their associated phenotypes; Cc: *Caulobacter crescentus*, Mx: *Myxococcus xanthus.* Refer to **Table S1** for a more detailed summary of mutant phenotypes.

## RESULTS

### Strategy for trapping and imaging McdA at specific steps of its ATPase cycle

We performed *in vivo* fluorescence microscopy to determine how McdA dynamics and carboxysome organization were altered for McdA mutants trapped at specific steps of its ATP cycle. To visualize carboxysomes, the fluorescent protein monomeric Turquoise2 (mTQ) was fused to the C-terminus of the small subunit of the Rubisco enzyme (RbcS) yielding RbcS-mTQ. RbcS-mTQ was expressed using a second copy of its native promoter (inserted at neutral site 1) in addition to wild-type *rbcS* at its native locus. To simultaneously image the McdA trap mutants in our carboxysome reporter strain, the amino acid substitutions were made in the ATP-binding pocket of an McdA variant that was N-terminally fused to the fluorescent protein monomeric NeonGreen (mNG) (Shaner *et al.*, 2013). We have previously shown that mNG-McdA is fully functional for carboxysome positioning when expressed as the only copy of McdA at its native locus (MacCready *et al.*, 2018). Finally, we also performed Phase Contrast imaging to monitor for changes in cell morphology, as we have recently shown that carboxysome mispositioning in *mcdA* or *mcdB* deletion strains triggers cell elongation, which we proposed is a response to carbon limitation (Rillema *et al.*, 2020).

### ATP-binding and dimerization mutants of McdA are diffuse in the cytoplasm and carboxysomes are mispositioned

We first set out to determine the *in vivo* localization pattern of McdA when unbound from ATP, and its impact on carboxysome positioning. We substituted the invariant catalytic Lysine to an Alanine (K15A) or Glutamine (K15Q) in the deviant Walker A box of McdA **(Figure 1B)**. Synonymous mutations in several other ParA-type ATPases have been shown to prevent ATP-binding **(Figure 1C)**. In wild-type *S. elongatus* cells, as shown previously, mNG-McdA oscillates on the nucleoid to equidistantly position RbcS-mTQ-labeled carboxysomes down the long axis of the cell **(Figure 2A)**. Both ATP-binding mutants of McdA no longer oscillated on the nucleoid, but rather were found to be diffuse in the cytoplasm and carboxysomes were mispositioned **(Figure 2, B–C)**. We then substituted the invariant Glycine to a Valine (G11V) in the deviant Walker A box of McdA **(see Figure 1, B–C)**, which allows for ATP-binding, but the bulky side-chain of Valine sterically prevents dimerization (Lutkenhaus, 2012). As with the ATP-binding mutants, the dimerization mutant of McdA was also diffuse in the cytoplasm, and carboxysomes were no longer uniformly distributed in the cell **(Figure 2D)**.

**Figure 2:**
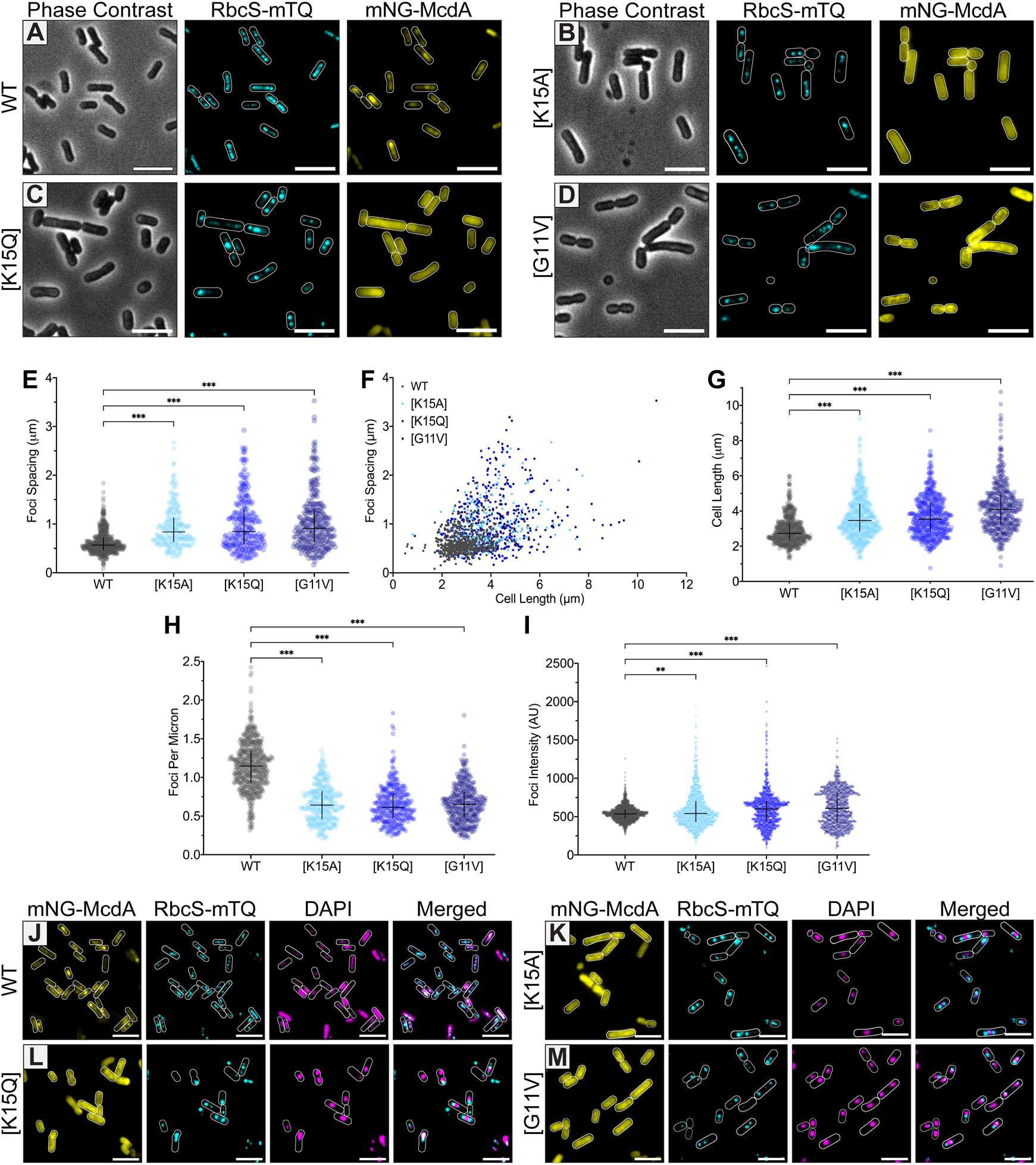
McdA mutants deficient in ATP binding and dimerization are unable to interact with the nucleoid and position carboxysomes. (**A**) mNG-McdA dynamically oscillates and positions carboxysomes labelled with RbcS-mTQ (cyan). (**B-D**) ATP-binding (K15A and K15Q) and dimerization (G11V) mutants of mNG-McdA no longer oscillate and carboxysomes aggregate. Cell outlines in fluorescent channels are based on the Phase Contrast image. (**E**) Spacing between carboxysome foci in the same cell. (**F**) Distribution of spacing between carboxysome foci as a function of cell length. For (**E**) and (**F**): WT *n* = 558 cells; *n* > 200 cells per mutant strain. (**G**) Cell lengths of specified strains. *n* > 400 cells per strain. (**H**) Number of carboxysome foci per unit cell length for each strain. WT *n* = 578 cells; *n* > 300 cells per mutant strain. (**I**) Carboxysome foci intensity for each cell strain (Arbitrary Units = AU). WT *n* = 1925 foci; n > 1100 foci per mutant strain. Data represent median with interquartile range. *** p < 0.001, ** p < 0.005 by Kruskal-Wallis test. (**J-M**) Microscopy images of cells with ciprofloxacin-compacted nucleoids. mNG-McdA and the specified variants (yellow), carboxysome foci (cyan) and DAPI-stained nucleoids (magenta). Carboxysome and DAPI channels are merged. Scale bars: 5 μm.

When we compared the nearest-neighbor spacing of carboxysome foci as a function of cell length, wild-type showed the same uniform spacing (0.6 ± 0.2 μm) regardless of cell length **(Figure 2, E–F)**. All three mutants, on the other hand, displayed increased spacing, and variability in spacing, as cell length increased **(Figure 2, E–F)**. The average cell lengths of the ATP-binding and dimerization mutants were significantly longer compared to wild-type **(Figure 2G)**; a change in cell morphology that mirrors the Δ*mcdA* phenotype **(Supplementary Figure S1A)** (Rillema *et al.*, 2020).

The increased spacing resulted in fewer carboxysome foci per unit cell length **(Figure 2H)**. Comparing the fluorescence intensity of carboxysome foci suggested that the increased spacing in all three mutant populations was the result of carboxysome aggregation **(Figure 2I)**. Overall, McdA mutant strains defective for ATP-binding and dimerization displayed a cell elongation phenotype, and possessed few and irregularly-spaced carboxysome aggregates. These phenotypes match what we have previously observed in the *mcdA* deletion strain (Rillema *et al.*, 2020), which suggests a complete loss of function in carboxysome positioning when McdA cannot bind ATP and dimerize.

### ATP-binding and dimerization are required for McdA to position carboxysomes on the nucleoid

Plasmids deleted for their ParA-type partitioning system are no longer distributed along the nucleoid. Rather, the plasmids become nucleoid ‘excluded’ (Erdmann *et al.*, 1999; Ringgaard *et al.*, 2009; Vecchiarelli *et al.*, 2012; Planchenault *et al.*, 2020). We have shown that nucleoid exclusion also occurs for carboxysomes in *S. elongatus* strains deleted for *mcdA* or *mcdB* (MacCready *et al.*, 2018). We set out to determine if carboxysomes are nucleoid excluded in the ATP-binding and dimerization mutants of McdA. Due to the polyploid nature of *S. elongatus*, DAPI staining does not easily resolve the nucleoid region from the cytoplasm **(Figure S1B)**. We therefore used the gyrase inhibitor ciprofloxacin to induce nucleoid compaction, which increased the cytoplasmic space observable by epifluorescence microscopy. Conveniently, when wild-type *S. elongatus* cells were treated with ciprofloxacin, mNG-McdA still oscillated on the compacted nucleoid **(Movie S1)**, and carboxysomes were still distributed over the nucleoid region of the cell and not in the cytoplasmic spaces **(Figure 2J)**. The ATP-binding and dimerization mutants of mNG-McdA, on the other hand, remained diffuse in the cytoplasm and carboxysomes were nucleoid excluded, but in a surprising manner **(Figure 2, K–M)**. Rather than having carboxysomes randomly distributed in the cytoplasmic region of the cell, the carboxysome aggregates butted-up against the ends of the compacted nucleoids **(Figure 2, K–M merged panels, and Figure S1C)**. A similar observation was recently found for plasmids lacking their partition system (Planchenault *et al.*, 2020), suggesting this is a widespread mesoscale phenomenon for both genetic and proteinaceous complexes in a bacterial cell.

Many ParA family ATPases are monomeric in their apo forms and dimerize upon ATP-binding, which then licenses non-specific DNA binding *in vitro* or nucleoid binding *in vivo* (Lutkenhaus, 2012; Kiekebusch and Thanbichler, 2014). Taken together, our data suggest that ATP-binding and dimerization are prerequisite steps needed for McdA to bind the nucleoid and distribute carboxysomes within the nucleoid region of the cell.

### The ATP-Trap mutant McdA[D39A] does not associate with the nucleoid or McdB *in vivo*

To solve the sandwich-dimer structure of an McdA homolog from the cyanobacterium *Cyanothece* sp. PCC 7424, the Schumacher group made an ATP-trap mutant by substituting the catalytic Aspartate residue to an Alanine in the Walker A’ box **(see Figure 1, A–B)** (Schumacher *et al.*, 2019). Synonymous ParA family mutants have been shown to form ATP-bound dimers competent for DNA-binding and interaction with their cognate ParB, but are deficient in ATP-hydrolysis **(see Figure 1C)**. We made the corresponding mutation in McdA (D39A) to determine the *in vivo* localization pattern of an McdA mutant presumably trapped as an ATP-bound dimer, and its effect on carboxysome positioning. Unexpectedly, mNG-McdA[D39A] was diffuse in the cytoplasm and carboxysomes were mispositioned in a manner that was identical to our ATP-binding and dimerization mutants of McdA **(Figure S2A)**. The data suggests McdA[D39A] cannot bind the nucleoid due to a loss in non-specific DNA binding activity. The Schumacher group showed that ATP-bound McdA[D38A] from *Cyanothece* can dimerize and bind a non-specific DNA substrate *in vitro* (Schumacher *et al.*, 2019), however the interaction affinity with DNA was not compared to wild-type McdA. Since *S. elongatus* McdA is highly insoluble, we purified the McdA homolog from *Cyanothece* (*Ct*McdA) and its ATP-trap variant *Ct*McdA[D38A] (used to solve the McdA structure), and found via Electrophoretic Mobility Shift Assays that *Ct*McdA[D38A] has significantly reduced DNA-binding activity compared to wild-type **(Figure S2B)**, which is consistent with our *in vivo* observations of the corresponding mutant in *S. elongatus* **(Figure S2A)**. We also performed Bacterial Two-Hybrid assays and found that while wild-type McdA showed a strong interaction with McdB, McdA[D39A] did not **(Figure S2C)**, which also explains our *in vivo* observations of this mutant in *S. elongatus* **(Figure S2A)**. We propose the ATP-trapped dimer of McdA[D39A] does not go through the conformational change that licenses nucleoid binding, which our data suggest is a prerequisite for McdB interaction and distributing carboxysomes over the nucleoid.

### The ATP-Trap mutant McdA[K15R] locks onto McdB-bound carboxysomes

We set out to construct another ATP-trap mutant of McdA that can adopt the nucleoid binding state and interact with McdB. Arguably the best studied ATP-trap mutant from the ParA family of ATPases comes from the P1 plasmid partitioning system (Fung *et al.*, 2001). Mutating the catalytic Lysine to an Arginine in the deviant Walker A box of P1 ParA (K122R) has shown robust *in vitro* and *in vivo* phenotypes **(see Figure 1, B–C)**. *In vitro*, ParA[K122R] can bind ATP, dimerize, and bind non-specific DNA with an affinity comparable to wild-type, but irreversibly associates with ParB because ParB cannot stimulate the ATPase activity required for releasing this association (Fung *et al.*, 2001; Vecchiarelli *et al.*, 2013). *In vivo*, ParA[K122R] results in a worse-than-null and dominant-negative phenotype called Par^PD^ for “propagation-defective”, whereby plasmids are less stable than when they have no partition system at all (Youngren and Austin, 1997). Given the severity of the mutation, the mechanism for the Par^PD^ phenotype has not been directly identified *in vivo*. However, the inability to disassemble the DNA-ParA[K122R]-ParB-plasmid complex *in vitro* suggests a likely mechanism (Hwang and Vecchiarelli *et al.*, 2013).

Strikingly, the corresponding ATP-trap mutant in mNG-McdA (K15R) resulted in nearly complete colocalization with carboxysomes **(Figure 3A)**. When *mcdB* was deleted from this strain, mNG-McdA[K15R] no longer associated with carboxysomes; instead coating the nucleoid thus showing its ability to still bind non-specific DNA **(Figure 3B)**. The data suggest that the ATP-trap mutant, McdA[K15R], locks carboxysomes onto the nucleoid via an irreversible interaction between McdA and McdB. Consistently, Bacterial-2-Hybrid analysis showed that McdA[K15R] associates more strongly with McdB compared to wild-type McdA **(Figure 3C)**, while all other McdA mutants studied thus far showed no interaction with McdB **(Figure S2C)**.

**Figure 3:**
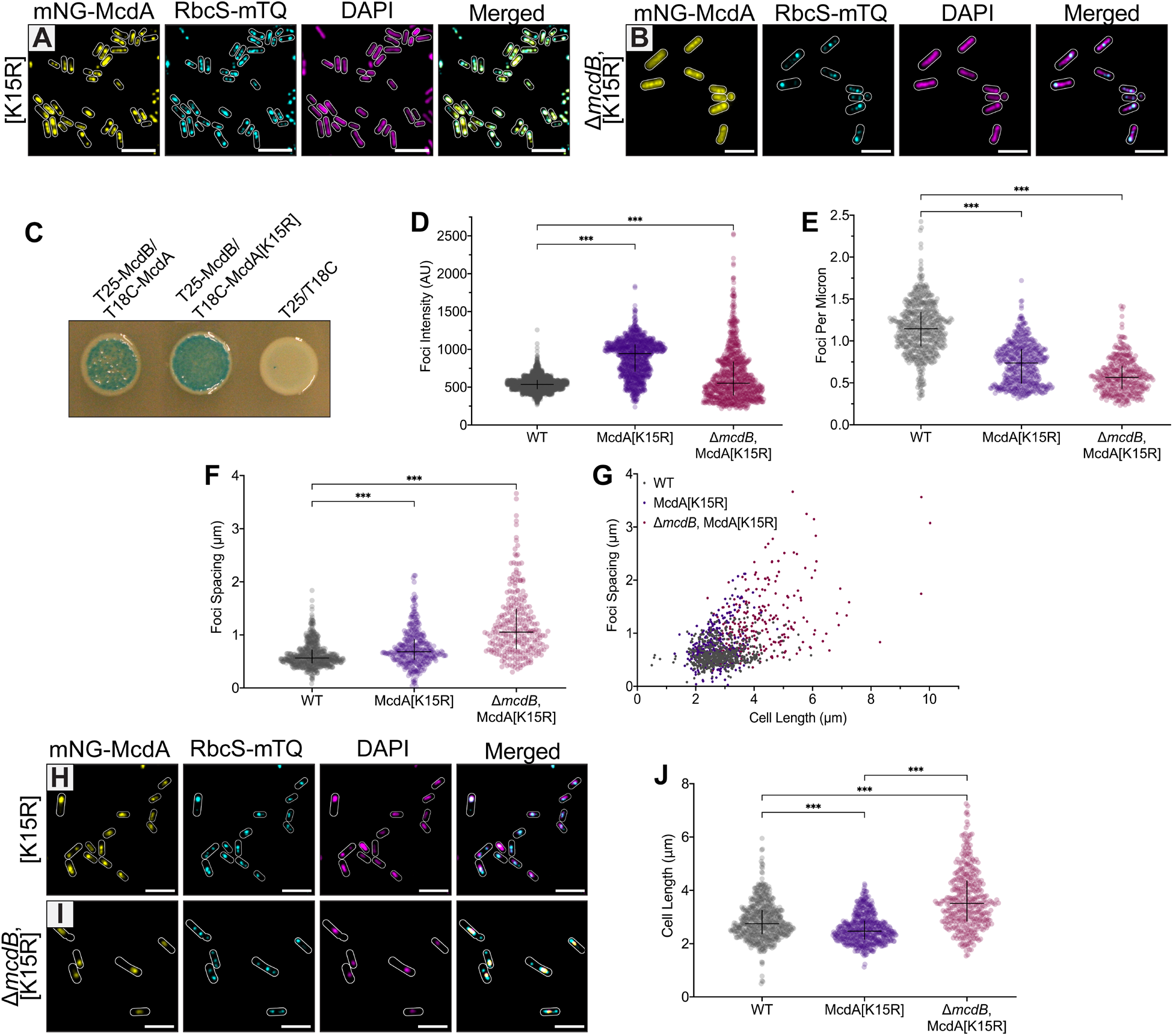
The ATP-Trap mutant McdA[K15R] irreversibly locks McdB-bound carboxysomes onto the nucleoid. (**A**) Microscopy images of mNG-McdA[K15R] (yellow), RbcS-mTQ-labelled carboxysomes (cyan) and DAPI-stained nucleoid (magenta). Merged image overlays mNG-McdA[K15R] and RbcS-mTQ-labelled carboxysomes. (**B**) Microscopy images of mNG-McdA[K15R] in *ΔmcdB* background strain. Merged image shows carboxysome and DAPI signals. (**C**) Bacterial two-hybrid interaction assay between the indicated protein pairs. Image is representative of three independent experiments. (**D**) Carboxysome foci intensity for specified cell strains. (Arbitrary Units = AU). WT *n* = 1925 foci; *n* > 800 foci per mutant strain. (**E**) Carboxysome foci number as a function of cell length. WT *n* = 578 cells; *n* > 370 cells of mutant strains. (**F**) Spacing of carboxysome foci. WT *n* = 558 cells; *n* > 250 cells of mutant strains. (**G**) Distribution of spacing between carboxysome foci as a function of cell length. WT *n* = 558 cells; *n* > 380 cells of mutant strains. (**H**) Microscopy images of mNG-McdA[K15R] (yellow), carboxysome foci (cyan) and DAPI-stained nucleoid (magenta) with ciprofloxacin treatment. Merged image shows carboxysome and DAPI signals. (**I**) Microscopy images of mNG-McdA[K15R] (yellow) in *ΔmcdB* background strain, carboxysome foci (cyan) and DAPI-stained nucleoid (magenta) with ciprofloxacin treatment. Merged image overlays carboxysome, mNG-McdA[K15R] and DAPI. (**J**) Cell lengths of specified strains. WT *n* = 558 cells; *n* > 250 cells of mutant strains. Scale bars: 5 μm.

Compared to wildtype, the McdA[K15R] mutant displayed significantly higher carboxysome foci intensities; a phenotype that was dependent on the presence of McdB **(Figure 3D)**. Consistent with carboxysome aggregation, the McdA[K15R] mutant displayed fewer carboxysome foci per unit cell length **(Figure 3E)**. The data suggest that McdB-stimulated ATP hydrolysis by McdA is required to disaggregate and distribute carboxysomes in the cell.

### The ATP-Trap mutant McdA[K15R] locks McdB-bound carboxysomes onto the nucleoid

Intriguingly, McdA[K15R] in the *mcdB* deletion strain displayed increased carboxysome spacing, and variability in spacing, as cell length increased **(Figure 3, F–G)**; a phenotype that is identical to an *mcdA* null mutant (Rillema *et al.*, 2020). With McdB present however, the McdA[K15R] strain had carboxysome spacing closer to that of wild-type **(Figure 3F)**. Unique to the McdA[K15R] mutant, carboxysome foci were enriched within the midcell region **(Figure S3)**. Also unlike all other McdA mutants described thus far, which were diffuse in the cytoplasm with nucleoid-excluded carboxysomes, mNG-McdA[K15R] strongly colocalized with carboxysomes over ciprofloxacin-compacted nucleoids **(Figure 3H)**. In the *ΔmcdB* background, mNG-McdA[K15R] remained associated with the compacted nucleoid, once again showing this mutant retains non-specific DNA binding activity, while carboxysomes became nucleoid excluded **(Figure 3I)**. Together, the data show that the ATP-trap mutant McdA[K15R] locks carboxysome aggregates onto the nucleoid via an irreversible interaction with McdB.

Finally, we asked if locking carboxysome aggregates onto the nucleoid in the McdA[K15R] strain resulted in the same cell elongation phenotype found for all other McdA mutants described thus far. Surprisingly, the McdA[K15R] strain did not elongate **(Figure 3J)**. In fact, the McdA[K15R] cells were slightly smaller than wild-type. When *mcdB* was deleted in the McdA[K15R] strain, the cell elongation phenotype returned. The findings suggest that the pseudo-positioning of carboxysome aggregates locked onto the nucleoid is sufficient to prevent cell elongation induced by the mispositioning of nucleoid-excluded carboxysome aggregates in null mutants of the McdAB system (Rillema *et al.*, 2020).

### McdA represents an unstudied subclass of ParA-family ATPases

Despite the McdA structure adopting an ATP sandwich dimer as shown for other ParA ATPases (Schumacher *et al.*, 2019), McdA lacks the classical “Signature Lysine” residue in the deviant Walker A box that defines this family **(see Figure 1B)**. Instead, the McdA structure identified a lysine residue, not only outside of the deviant Walker A box, but in the C-terminal half of the protein at position 151, which is employed as the Signature Lysine **(Figure 4A)** (Schumacher *et al.*, 2019). As with the classical signature Lysine, Lys151 interacts with the ATP molecule bound in the adjacent McdA monomer; making the same cross contacts to the oxygen atom connecting the β- and γ-phosphates. Sequence alignments of McdA homologs that lack the classical signature Lysine in the deviant Walker A box, invariably encode for a lysine that corresponds to Lys151 in *S. elongatus* McdA **(Figure 4A)**. Given the McdA structure, sequence conservation, and biochemical data suggesting Lys151 is important for ATP binding and dimerization, we next observed the effect of mutating Lys151 to an Alanine *in vivo*. The majority of mNG-McdA[K151A] remained diffuse in the cytoplasm, while a minor fraction colocalized with few and irregularly spaced carboxysome aggregates **(Figure 4B)**. Carboxysome foci intensity, spacing and average cell length were identical to that found for the other ATP-binding and dimerization mutants of McdA tested in this study **(Figure S4, A–D)**. Also, ciprofloxacin treatment showed carboxysome aggregates were nucleoid excluded, and once again butted-up against the nucleoid poles **(Figure 4C)**. The findings highlight the importance of Lys151 as the “Signature Lysine” for an unstudied ParA subclass, in forming the ATP-bound McdA dimer competent for nucleoid binding and positioning carboxysomes.

**Figure 4:**
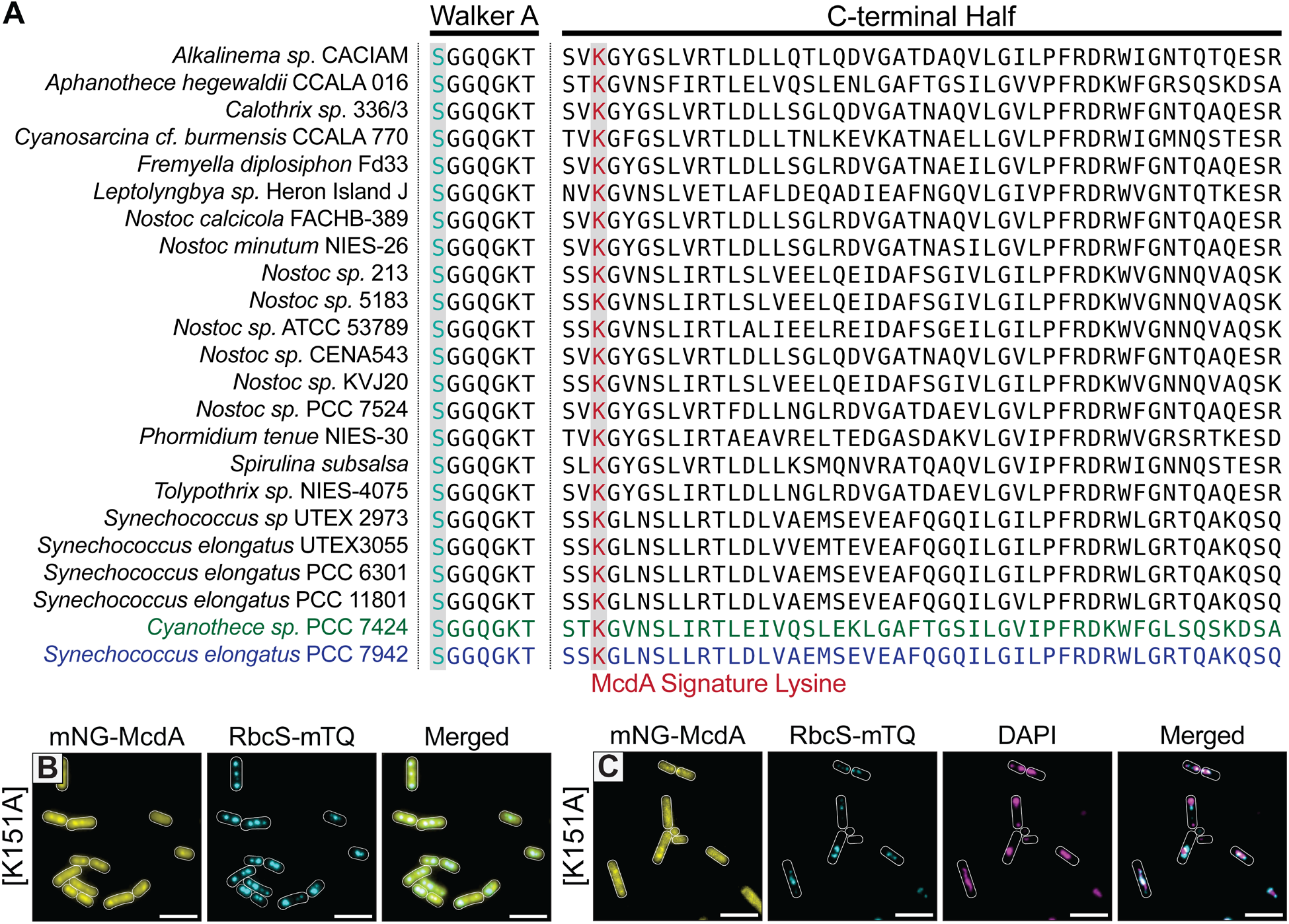
McdA is a member of an unstudied subclass of ParA-type ATPase characterized by a different signature lysine position. (**A**) Sequence alignment of McdA homologs possessing a serine residue in place of the signature lysine the Walker A box that co-occurs with an invariant lysine residue in the C-terminal half of proteins – the McdA signature lysine. (**B**) Microscopy images of mNG-McdA[K151A] and RbcS-mTQ-labelled carboxysomes (cyan). (**C**) Microscopy images of mNG-McdA[K151A] (yellow), carboxysome foci (cyan) and DAPI-stained nucleoid (magenta) with ciprofloxacin treatment. Merged image shows carboxysome and DAPI signals. Scale bars: 5 μm.

### Moving the Signature Lysine of McdA into the Walker A box reconstitutes carboxysome pseudo-positioning

Remarkably, Lys151 of the McdA structure overlays exceptionally well onto the signature lysine position in the deviant Walker-A box of classical ParA family members (Schumacher *et al.*, 2019). This finding suggested that it may be possible to maintain carboxysome positioning with an McdA mutant that has its signature Lysine at position 151 reintroduced into the classical position in the deviant Walker A box at position 10 **(see Figure 4A)**. To make the signature Lysine mutant, McdA[S10K, K151S], we swapped the Serine at position 10 in the deviant Walker A box with the Lysine at position 151. The mNG-McdA[K151S] phenotype mirrored that of McdA[K151A] - largely diffuse in the cytoplasm with nucleoid-excluded carboxysome aggregates **(Figure S4E)**. mNG-McdA[S10K, K151S], on the other hand, largely colocalized with carboxysome foci **(Figure 5A)**. Also, carboxysome spacing **(Figure 5B)** and intensity **(Figure 5C)** both trended back towards wild-type values, and ciprofloxacin treatment showed that carboxysomes were now positioned within the nucleoid region of the cell **(Figure 5D)**. Together, the data suggest a pseudo-restoration of carboxysome positioning on the nucleoid. Consistently, the McdA[S10K, K151S] cell population had cell lengths revert back to wild-type **(Figure 5E)**, suggesting this pseudo-positioning of carboxysomes is sufficient to alleviate the cell elongation mutant phenotype (Rillema *et al.*, 2020).

**Figure 5:**
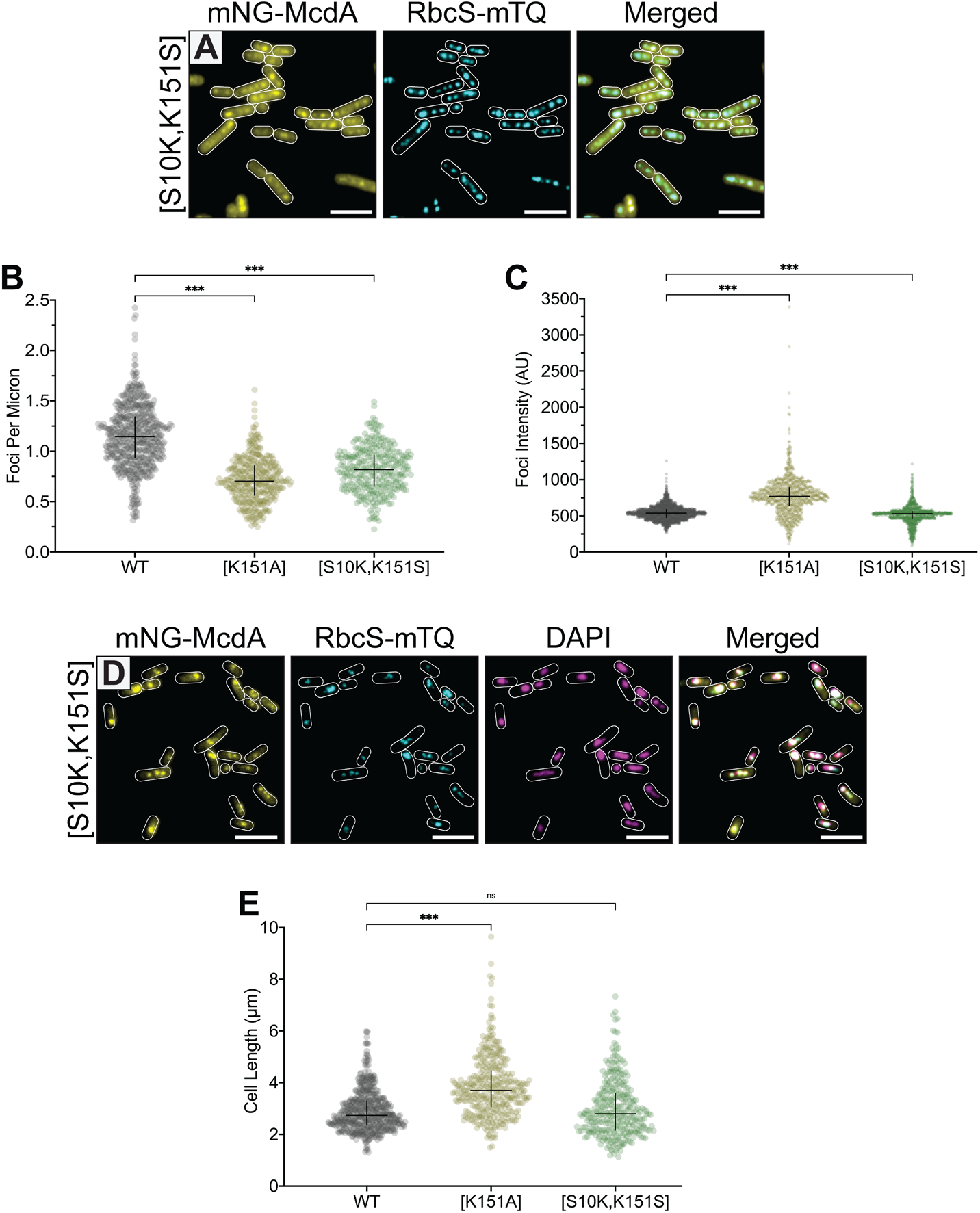
Carboxysomes are pseudo-positioned when the McdA signature lysine is moved into the classical Walker A box position. (**A**) Microscopy images of mNG-McdA[S10K, K151S] and RbcS-mTQ-labelled carboxysomes (cyan). (**B**) Number of carboxysome foci per unit cell length. WT *n* = 578 cells; *n* > 350 cells of mutant strains. (**C**) Carboxysome foci intensity. (Arbitrary Units = AU). WT *n* = 1925 foci; *n* > 950 foci from mutant strains. (**D**) Microscopy images of mNG-McdA[S10K, K151S] (yellow), carboxysome foci (cyan) and DAPI-stained nucleoid (magenta) with ciprofloxacin treatment. Merged image overlays carboxysome, mNG-McdA[S10K, K151S] and DAPI signals. (**E**) Cell lengths of specified strains. ns = not significant by Kruskal-Wallis test. WT *n* = 561 cells; *n* > 320 cells of mutant strains. Scale bars: 5 μm.

## DISCUSSION

Members of the ParA family of ATPases position a wide variety of genetic and proteinaceous cargos involved in diverse biological processes (Lutkenhaus, 2012; Vecchiarelli *et al.*, 2012; Kiekebusch and Thanbichler, 2014). ATP cycling by the ParA ATPase is critical for its dynamic patterning behavior in the cell as well as its positioning activity on the cognate cargo. We recently found that the McdAB system is widespread across cyanobacteria and carboxysome-containing proteobacteria (MacCready *et al.* 2020; MacCready and Tran *et al.* 2021), yet it remains unknown how the ATPase cycle of McdA controls its oscillatory dynamics and its function in distributing carboxysomes across the nucleoid length. Several well-researched amino acid substitutions in the conserved ATP-binding site of ParA family ATPases have been used to trap the ATP cycle at specific steps. These trap mutants have served as useful probes for dissecting the molecular steps involved in ParA-based positioning reactions **(Summary in Figure 1C, detailed in Table S1)**. To dissect how ATP mediates McdA function in positioning fluorescently-labelled carboxysomes, we introduced synonymous amino acid substitutions in the ATP-binding pocket of fluorescently-labelled McdA to trap it at specific steps of the ATP cycle. The phenotypes of these trap mutants have allowed us to correlate the known biochemistry of well-studied ParA family ATPases with specific steps in McdA action we observed here *in vivo*.

Overall we find that ATP-binding, dimerization, and an ATP-specific conformational change in McdA are all prerequisite steps for McdA to associate with the nucleoid via non-specific DNA binding activity **(Figure 6A)**. Our findings suggest that McdB-bound carboxysomes can only interact with McdA in this DNA-binding state. Nucleoid-associated McdA tethers McdB-bound carboxysomes to the nucleoid. But ultimately, McdB stimulates ATP-hydrolysis by McdA, which reverts McdA back into its monomeric form that can no longer bind the nucleoid in the vicinity of the carboxysome. Through this Brownian-ratchet mechanism (MacCready *et al.*, 2018), McdB-bound carboxysomes are uniformly distributed as they locally generate McdA depletion zones on the nucleoid, and then move up the resulting McdA gradient towards higher concentrations **(Figure 6B)**.

**Figure 6:**
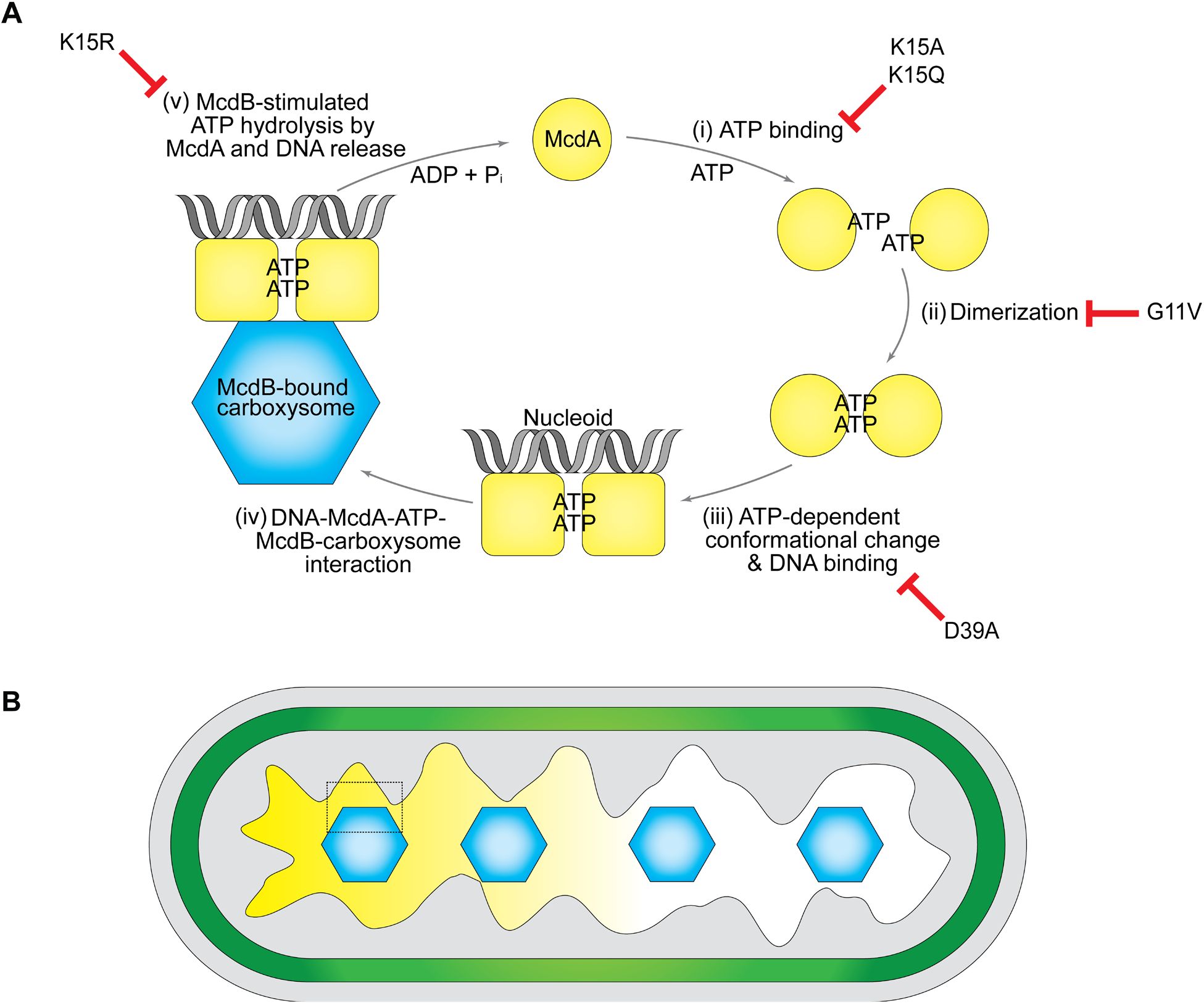
Model for ATP-cycling by McdA and associated functions in carboxysome positioning. (**A**) The ATPase cycle of McdA. Trap mutants of McdA identified in this study are indicated. (*i*) When unbound from ATP, McdA monomers are diffuse in the cytoplasm. (*ii*) Upon ATP-binding, McdA is competent for dimerization. (*iii*) ATP-bound McdA dimers must go through an ATP-dependent conformational change that licenses non-specific DNA binding to the nucleoid. (*iv*) McdB-bound carboxysomes are tethered via interactions with McdA-ATP dimers on the nucleoid. (*v*) McdB stimulates McdA ATPase activity and its release from the nucleoid in the vicinity of a carboxysome. (**B**) McdB-bound carboxysomes are uniformly distributed as they continually move toward higher concentrations of McdA on the nucleoid. The dashed box indicates the cellular region magnified in (A).

McdA mutants unable to bind ATP, dimerize, or undergo the ATP-specific conformational change required for nucleoid binding were diffuse in the cytoplasm and carboxysomes were observed as nucleoid-excluded aggregates. These mutant strains also displayed cell elongation. We have recently shown that *mcdA* and *mcdB* deletion strains also elongate (Rillema *et al.*, 2020). Heterotrophic bacteria have been shown to undergo cell elongation as a carbon-limitation response (Rangarajan *et al.*, 2020). We also recently proposed that carboxysome aggregation results in decreased carbon-fixation efficiency, and that cell elongation is a response triggered by the resulting carbon limitation in this photoautotroph (Rillema *et al.*, 2020). Since the phenotype of these McdA trap mutants mirror the *mcdA* deletion strain, our findings suggest a complete loss of function in carboxysome positioning when McdA cannot bind ATP, dimerize, and adopt its nucleoid-binding conformation.

### Nucleoid-excluded carboxysomes are trapped at the cytoplasm-nucleoid interface

We previously showed that in Δ*mcdA* or Δ*mcdB* strains of *S. elongatus*, carboxysomes still fully assemble, but coalesce into nucleoid-excluded aggregates (MacCready *et al.*, 2018). Given the polyploid nature of *S. elongatus*, there is insufficient cytoplasmic space to resolve whether carboxysomes aggregated due to physical interactions with each other, or if they simply coalesced because of nucleoid exclusion. We used the gyrase-inhibitor ciprofloxacin to compact the nucleoid and increase the cytoplasmic space of *S. elongatus* cells. Surprisingly, we found that in the absence of a functional McdAB system, carboxysome aggregates did not diffuse into the increased cytoplasmic space of ciprofloxacin-treated cells. Instead, the aggregates were maintained at the cytoplasm-nucleoid interface. It was recently shown that large plasmids lacking their ParA-based partition system, or large DNA circles excised from the chromosome, also localize to this interface (Planchenault *et al.*, 2020). This phenomenon was plasmid-size dependent; only plasmids larger than 100 kb preferentially localized to the nucleoid edge and did not diffuse into the nucleoid-free cytoplasmic space of the cell. Our findings here show that this preferential localization to the nucleoid edge is not specific to plasmids, but is rather a widespread phenomenon in bacteria for both genetic and proteinaceous complexes on the mesoscale. Given the size-dependence of nucleoid-evicted complexes being unable to penetrate the cytoplasm, we believe the most parsimonious explanation is that carboxysomes, and other mesoscale complexes, perceive the cytoplasmic environment as glassy (Parry *et al.*, 2014); thus exhibit caging and subdiffusive behaviors at the nucleoid-cytoplasm interface.

Remarkably, wildtype cells treated with ciprofloxacin still displayed mNG-McdA oscillations and carboxysomes were still distributed over the highly compacted nucleoid. The data suggest that the McdAB system can distribute carboxysomes regardless of whether the nucleoid is expanded or in an extremely compacted state. This finding has implications for identifying the forces responsible for carboxysome movement and positioning within the nucleoid region of the cell.

### The ATP-trap mutant McdA[K15R] locks carboxysomes onto the nucleoid

We identified the ATP-trap mutant, McdA[K15R], that locks the nucleoid-McdA-McdB-carboxysome ternary complex. In the absence of McdB, mNG-McdA[K15R] still coated the nucleoid showing that it retains non-specific DNA binding activity, but carboxysomes were nucleoid excluded. In the presence of McdB, mNG-McdA[K15R] completely colocalized with massive carboxysome aggregates over the nucleoid. Together the findings show that McdA on the nucleoid transiently interacts with McdB on carboxysomes. McdB then stimulates McdA ATP-hydrolysis and release from the nucleoid in the vicinity of carboxysomes, which allows for continued movement up the resulting McdA gradient. Without the ability to hydrolyze ATP, McdA[K15R] irreversibly associates with McdB and statically tethers carboxysomes to the nucleoid. Since the ATP cycle cannot rest, McdB-bound carboxysomes act as a sink for all McdA[K15R] in the cell, which explains the absence of mNG-McdA[K15R] redistribution across the nucleoid.

All McdA mutants that result in nucleoid-excluded carboxysome aggregation also showed a cell elongation phenotype. The McdA[K15R] strain, on the other hand, displayed carboxysome aggregates on the nucleoid and no cell elongation phenotype. In contrast, the cells were slightly shorter than wild-type. We have two hypotheses that could explain this phenotype. First, tethering carboxysomes to the nucleoid could allow for pseudo-positioning of carboxysomes. This “pilot-fish” mode of carboxysome positioning and inheritance could sufficiently improve carbon-fixation efficiency, thereby preventing carbon limitation and the cell elongation response. Alternatively, it can be envisioned that irreversibly tethering massive carboxysomes onto the nucleoid can have detrimental effects to a variety of DNA transactions such as DNA replication, transcription, nucleoid organization and compaction, and faithful chromosome segregation. Therefore the shorter cell length could simply be attributed to a slower growth rate.

### Swapping the Signature Lysine position in McdA resulted in carboxysome pseudo-positioning on the nucleoid

McdA represents a previously unstudied subclass of the ParA family, where the signature Lysine residue that defines this ATPase family is located in the C-terminal half of the protein, rather than in the Walker A box **(see Figure 4A)**. We find here that Lysine151 is indeed necessary for McdA to bind the nucleoid and position carboxysomes. Strikingly, we also found that repositioning this Lysine into the classical signature Lysine position in the Walker A box reconstituted carboxysome pseudo-positioning –carboxysome spacing and focal intensity trended back to wild-type values. This mutant also reverted back to wildtype cell lengths. However, the oscillatory dynamics observed with wildtype McdA were not reconstituted. Instead, mNG-McdA[S10K, K151S] colocalized with carboxysomes over the nucleoid. This mode of carboxysome positioning is similar to that observed for the P1 plasmid partition system. P1 ParB forms punctate foci by loading onto and around a DNA binding site called *parS* on the plasmid to be partitioned (Erdmann *et al.*, 1999; Sengupta *et al.*, 2010). The ParA ATPase uniformly distributes over the nucleoid, but also forms foci that colocalize with relatively immobile ParB-bound plasmids (Hatano and Niki, 2010). During plasmid partitioning and movement, the colocalized ParA foci disappear and only reappear once the sister plasmids have reached the ¼ and ¾ positions of the cell where they once again become relatively immobile. McdA has an ATPase activity two-orders of magnitude greater than ParA ATPases with a classical signature Lysine (MacCready *et al.*, 2018). It is attractive to speculate that the Lysine-swap mutant of McdA decreases its voracious ATPase activity, causing it to remain associated with McdB-bound carboxysomes for a longer period of time and adopting a “stick-and-move” mode of carboxysomes positioning over the nucleoid; similar to the P1 plasmid partition reaction described above.

Why does McdA have such a greater ATPase rate compared to classical ParA-type ATPases? We believe the answer lies in the difference in cargo copy-number in the cell. ParA-based DNA segregation systems are typically found on bacterial chromosomes and large low-copy plasmids. In both cases, the DNA is replicated and the sister copies are then segregated to opposing halves of the cell prior to division. Carboxysome copy number, on the other hand, can be significantly higher and varies depending on growth conditions. For example, when grown with high-light intensity, a single *S. elongatus* cell can contain up to a dozen carboxysomes (Sun *et al.*, 2016). We propose that for high-copy-number cargos, an increased ATPase activity is required to compensate for the decreased nearest-neighbor distance between adjacent cargos sharing the same nucleoid matrix. The increased ATPase rate would make the McdA gradient on the nucleoid more sensitive to carboxysome movements over these smaller spatial scales.

## MATERIALS AND METHODS

### Construct design

All constructs were made using Gibson assembly (Gibson *et al.*, 2009) from PCR fragments or synthesized dsDNA (Integrated DNA Technologies) and verified by Sanger sequencing. For *mcdB* deletion and native fluorescent fusion gene insertions into the *S. elongatus* genome, constructs were made as previously described (MacCready *et al.*, 2018).

### Growth conditions and transformations

All *S. elongatus* (ATCC^®^ 33912^™^) strains were grown in 125 mL baffled flasks (Corning) in 50 mL BG-11 medium (Sigma) pH 8.3 buffered with 1g/L HEPES. Cells were cultured in a Minitron incubation system (Infors-HT) with the following growth conditions: 60 μmol m^−2^ s^−1^ continuous LED 5600K light, 32°C, 2% CO_2_, and shaking at 130 RPM. Plasmids were cloned in chemically competent One Shot^™^ TOP10 *E. coli* cells (Thermo Fisher Scientific) in standard manipulation and culture conditions (Green and Sambrook 2012). Transformations of *S. elongatus* cells were performed as previously described (Clerico *et al.*, 2007). Transformant cells were plated on BG-11 agar with 12.5 μg/ml kanamycin, 12.5 μg/ml chloramphenicol or 25 μg/ml spectinomycin. Single colonies were picked and transferred into 96-well plates containing BG-11 medium with corresponding antibiotic concentrations. Complete gene insertions and absence of the wildtype gene were verified via PCR and cultures were removed from antibiotic selection by three series of back dilution prior to imaging.

### Ciprofloxacin treatment and nucleoid visualization

To induce nucleoid compaction, *S. elongatus* cells were incubated with 50 μM ciprofloxacin overnight under normal growth conditions. To visualize the compacted nucleoid region, ciprofloxacin-treated *S.elongatus* cells were harvested by centrifugation at 4,000 × g for 1 minute. The pelleted cells were then washed and resuspended in 100 μl of PBS (pH 7.2). DAPI (8 μl from a 20 μg/ml stock concentration) was added to the cell suspension followed by 20-minute incubation in the dark at 30°C. DAPI-stained cells were washed twice with 1 ml H_2_O, and then resuspended in 100 μl H_2_O prior to visualization using the DAPI channel.

### Fluorescence Microscopy

Exponentially growing cells (2 mls of cells at OD_750_ ~ 0.7) were harvested and spun down at 4,000 × g for 1 min, resuspended in 200 μl fresh BG-11 and 2 μl was then transferred to a 1.5% UltraPure agarose (Invitrogen) + BG-11 square pad on a glass-bottom dish (MatTek Life Sciences). All fluorescence and phase contrast imaging were performed using a Nikon Ti2-E motorized inverted microscope controlled by NIS Elements software with a SOLA 365 LED light source, a 100X Objective lens (Oil CFI Plan Apochromat DM Lambda Series for Phase Contrast), and a Photometrics Prime 95B Back-illuminated sCMOS camera. mNG-McdA variants were imaged using a “YFP” filter set (C-FL YFP, Hard Coat, High Signal-to-Noise, Zero Shift, Excitation: 500/20nm [490-510nm], Emission: 535/30nm [520-550nm], DichroicMirror: 515nm). RbcS-mTQ labelled carboxysomes were imaged using a “CFP” filter set (C-FL CFP, Hard Coat, High Signal-to-Noise, Zero Shift, Excitation: 436/20nm [426-446nm], Emission: 480/40nm [460-500nm], Dichroic Mirror: 455nm). DAPI fluorescence was imaged using a standard “DAPI” filter set (C-FL DAPI, Hard Coat, High Signal-to-Noise, Zero Shift, Excitation: 350/50nm [325-375nm], Emission: 460/50nm [435-485nm], Dichroic Mirror: 400nm). Image analysis was performed using Fiji v1.53b (Schindelin *et al.*, 2012).

### Image Analysis

Image analysis including cell segmentation, quantification of cell length, foci number, intensity and spacing were performed using Fiji plugin MicrobeJ 5.13I (Ducret *et al.*, 2016). Cell perimeter detection and segmentation were done using the rod-shaped descriptor with default threshold settings. Carboxysome detection was performed using the smoothed foci function with tolerance of 50 and Z-score of 30. Data were exported, further tabulated, graphed and analyzed using GraphPad Prism 9.0.1 for macOS, GraphPad Software, San Diego, California USA, https://www.graphpad.com.

### Bacterial Two-Hybrid

N -terminal T18 and T25 fusions of McdA, all McdA mutant variants, and McdB were constructed using the plasmids pKT25 and pUT18C. Plasmids were sequence-verified and co-transformed into *E. coli* BTH101 in both pairwise combinations (Karimova *et al.*, 1998). Several colonies of T18/T25 cotransformants were cultured in LB medium with 100 mg/ml ampicillin, 50 mg/ml kanamycin and 0.5 mM IPTG overnight at 30°C with 225 rpm shaking. Overnight cultures were spotted on indicator LB X-gal plates supplemented with 100 mg/ml ampicillin, 50 mg/ml kanamycin and 0.5 mM IPTG. Plates were incubated in the dark at 30°C up to 48 hours before imaging.

### Expression and purification of *Ct* McdA and *Ct* McdA[D38A]

Both *Ct*McdA and *Ct*McdA[D38A] were expressed and purified in a similar manner. For protein production, the expression plasmids for these constructs (Schumacher *et al.*, 2019) were transformed into *E. coli* C41(DE3) cells (Lucigen). Transformants were grown at 37°C and 225 rpm until an OD_600_ of 0.4–0.6 was reached. The culture flasks were rapidly cooled down to 15°C on and protein expression was then induced with the addition of 1 mM IPTG. After overnight induction, the cells were pelleted, flash frozen in liquid nitrogen and stored at −80°C. Harvested cells were resuspended in Buffer A (25 mM Tris-HCl pH 7.5, 300 mM NaCl, 10% glycerol, 0.5 mM BME, 50 mg/ml lysozyme, 1.25 kU benzonase, 2 Protease Inhibitor Cocktail tablets) and lysed using a probe sonicator with 15 s on, 15 s off pulsation for 8 min. The lysate was cleared by centrifugation at 12,000 × g at 4 °C for 40 min in a Fiberlite TM F15-8 × 50 cy Fixed Angle Rotor (ThermoFisher Scientific). The resulting lysate was filtered through a 0.45 μm syringe filter and loaded onto a 5 ml HiTrap^™^ TALON Crude cassette (GE) and eluted with a 0 to 400 mM imidazole gradient. Peak fractions were pooled and concentrated using an Amicon Ultra Centrifugal Device (10 KD MWCO). The concentrated protein sample was passed through a HiPrep 26/10 Desalting Column (GE) equilibrated in Q-Buffer (25mM Tris-HCl pH 7.5, 150 mM NaCl, 10% glycerol, 1mM EDTA, 1mM DTT). The sample was then immediately loaded onto a HiTrap^™^ Q HP 5 ml cassette (GE) equilibrated in Q-Buffer. The protein was eluted with a 150 mM to 2 M NaCl gradient. Peak fractions were concentrated to no more than 70 mM and flash frozen aliquots were kept at −80°C.

### DNA binding assay

Electrophoretic mobility shift assays (EMSAs) were performed in a final reaction volume of 10μl in a buffer containing 50 mM HEPES (pH 7.6), 5 mM MgCl_2_, and 100 mM KCl with 10nM pUC19 plasmid (2.8 kb) as the supercoiled DNA substrate. At the concentrations indicated, His-*Ct*McdA and His-*Ct*McdA[D38A] were incubated for 30 min at 23°C with or without ATP (1 mM). Reactions were then mixed with 1 μl 80 % glycerol, run on 1 % agarose gel in 1X TAE at 110V for 45 min and stained with ethidium bromide for imaging.

### Protein structure visualization and prediction

Molecular graphics and analyses of protein structures were performed with USCF Chimera, developed by the Resource for Biocomputing, Visualization and Informatics at the University of California, San Francisco, with support from NIH P41-GM103311 (Pettersen *et al.*, 2004). Prediction of *Se*McdA structure was performed with Phyre2 (Kelley *et al.*, 2015).

## ACKNOWLEDGEMENTS

We would like to thank Joshua MacCready, Kiyoshi Mizuuchi, Maria Schumacher and David Savage for helpful discussions. The pET15b expression vectors used for *Ct*McdA and *Ct*McdA[D38A] were kind gifts from Maria Schumacher. This work is supported by the National Science Foundation to A.G.V. (Award No. 1817478 and CAREER Award No. 1941966), Rackham Graduate Research Grant to P.H., Rackham Professional Development Grant to P.H., American Society for Microbiology Research Capstone Fellowship to P.H., Margaret Dow Towsley Scholarship to P.H. and by research initiation fund to A.G.V. provided by the MCDB department, University of Michigan.

## Abbreviations

BMC: Bacterial Microcompartment
mNG: monomeric NeonGreen
mTQ: monomeric Turquoise2
McdA: Maintenance of Carboxysome Distribution protein A
McdB: Maintenance of Carboxysome Distribution protein B
ParA: Partition protein A
ParB: Partition protein B
ATP: Adenosine triphosphate
AMPPNP: Adenylyl-imidodiphosphate
nsDNA: non-specific DNA
DAPI: 4′,6-diamidino-2-phenylindole

## Supplemental Information

### This File Contains

- Supplemental Figures S1 to S4
- Supplemental Movie Legend S1
- Supplemental Tables S1 and S2

---

**Figure S1:**
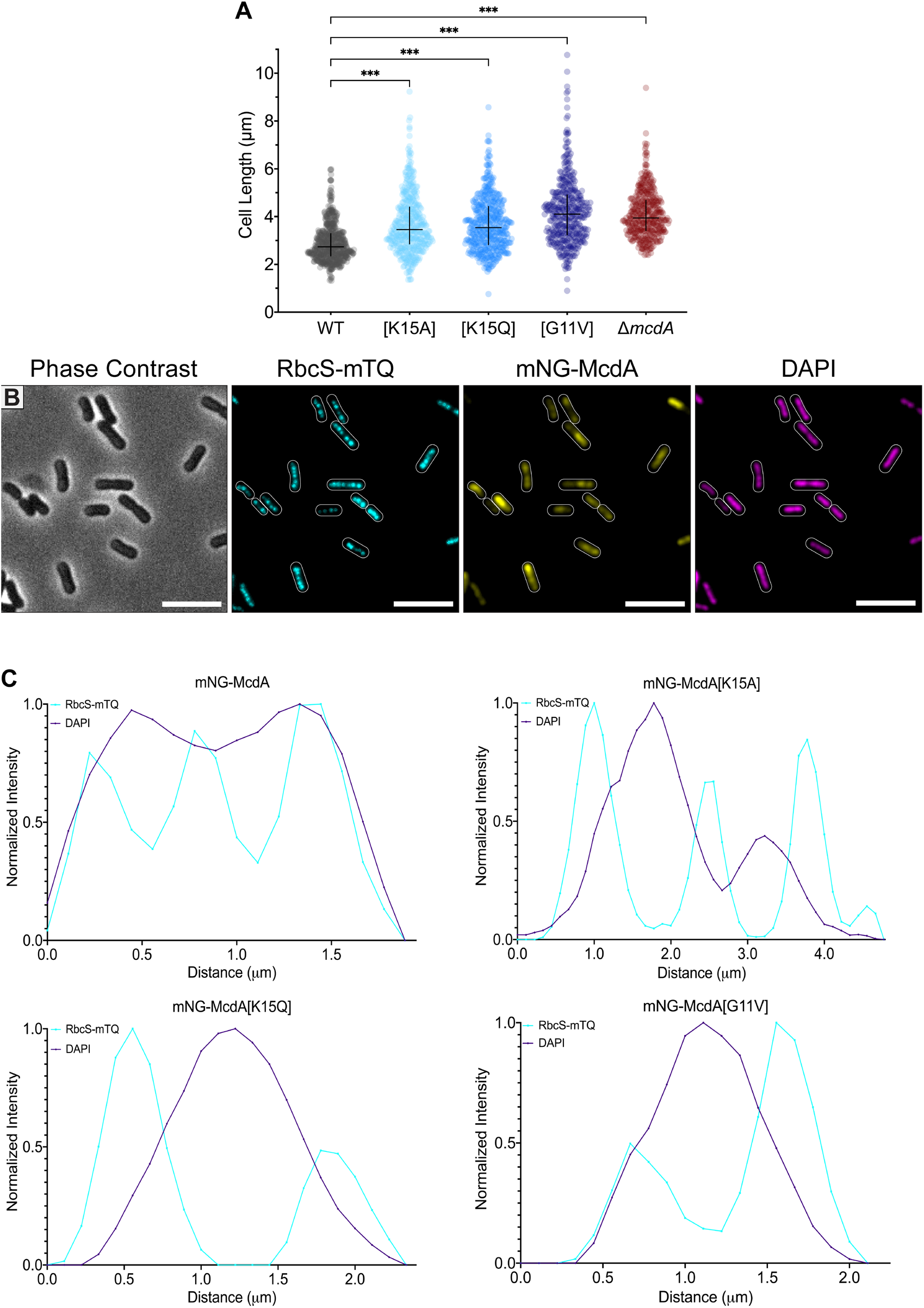
**(A)** Comparison of cell lengths among WT and specified mNG-McdA mutants and Δ*mcdA* strains. WT *n* = 558 cells; n > 380 cells per mutant strains. **(B)** Microscopy images of mNG-McdA cells (Figure 2A) with DAPI-stained nucleoid (magenta). **(C)** Line scans of carboxysome and nucleoid signals of specified strains. Each line scan graph is a representative signal measurement of cells from each strain.

**Figure S2:**
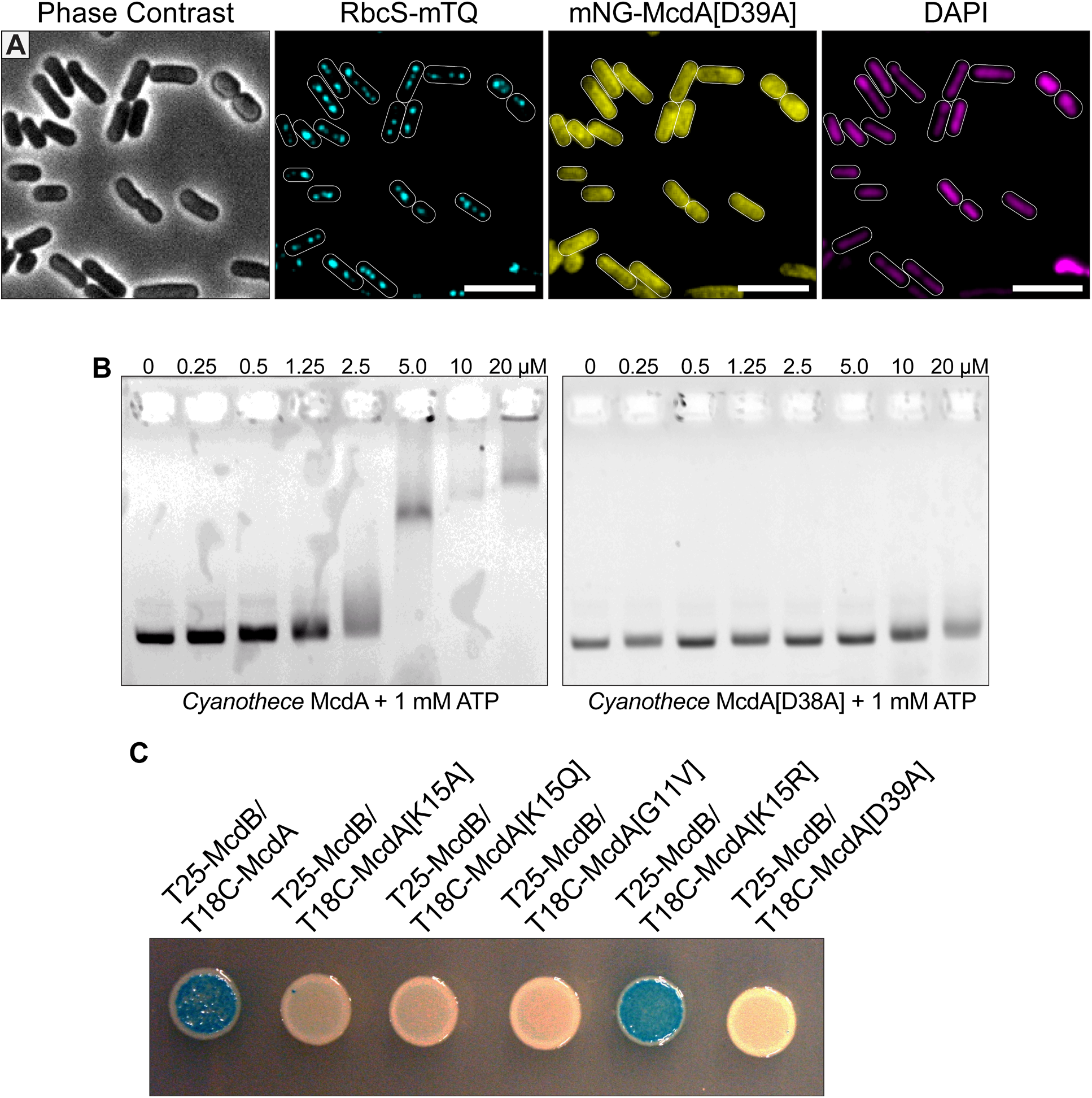
**(A)** Microscopy images of mNG-McdA[D39A] strain. **(B)** Bacterial two-hybrid interaction assay of McdB against the wild-type McdA or the specified mutants. Image is representative of three independent experiments. **(C)** Electrophoretic Mobility Shift Assay (EMSA) showing that wildtype *Ct*McdA binds and slows the migration of a non-specific plasmid DNA substrate in the presence of 1 mM ATP while *Ct*McdA[D38A] does not.

**Figure S3:**
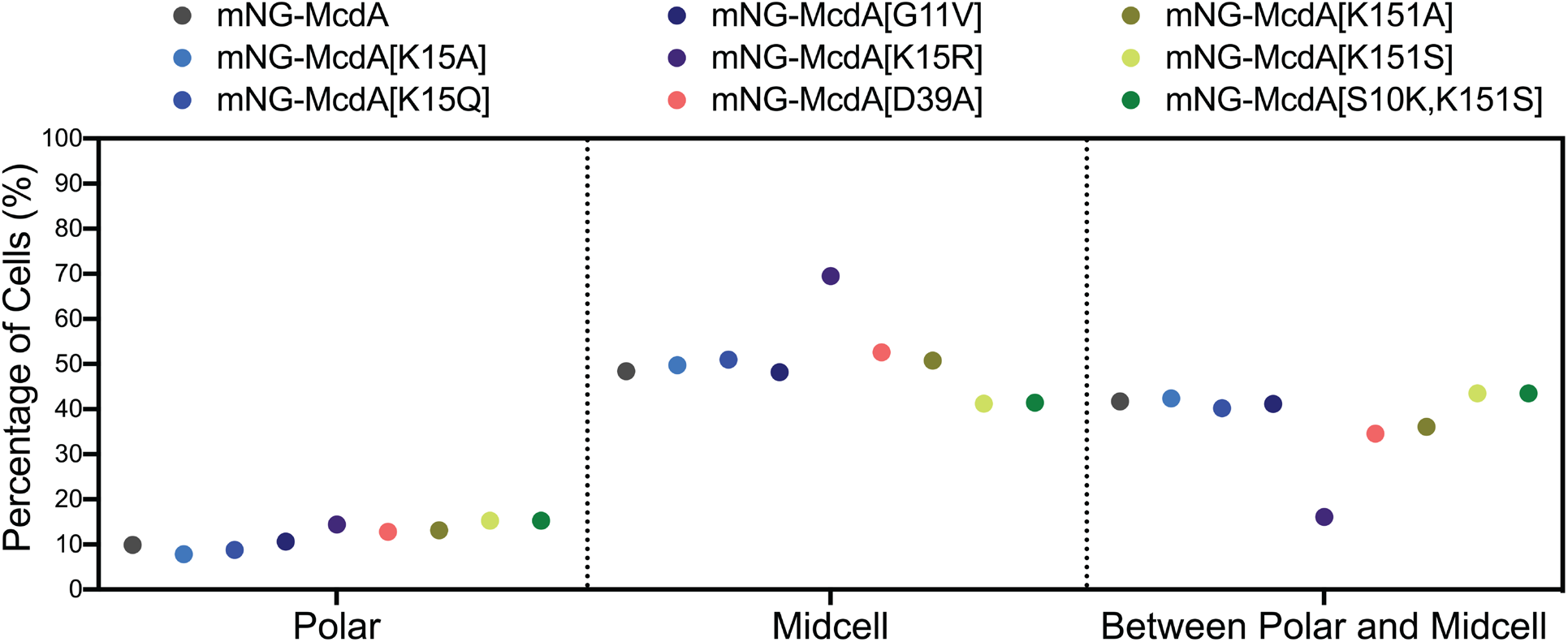
Binned subcellular localization of carboxysomes in the specified cell strains. Quantification was performed in MicrobeJ where carboxysome signals located within the region extending from the tip of the cell pole to a position on the medial axis located half the width away from the cell pole tip, are considered as “polar” localized. Carboxysome signals located within the region extending from the cell center to a position on the medial axis located half the width away from the cell center, are considered as “midcell” localized. Carboxysomes located between these two defined regions were grouped as “between polar and midcell”. The McdA[K15R] cell population significantly deviants from all other McdA variants in regard to carboxysome foci positioning in the cell. WT *n* = 1000 foci; n > 800 foci per mutant strain.

**Figure S4:**
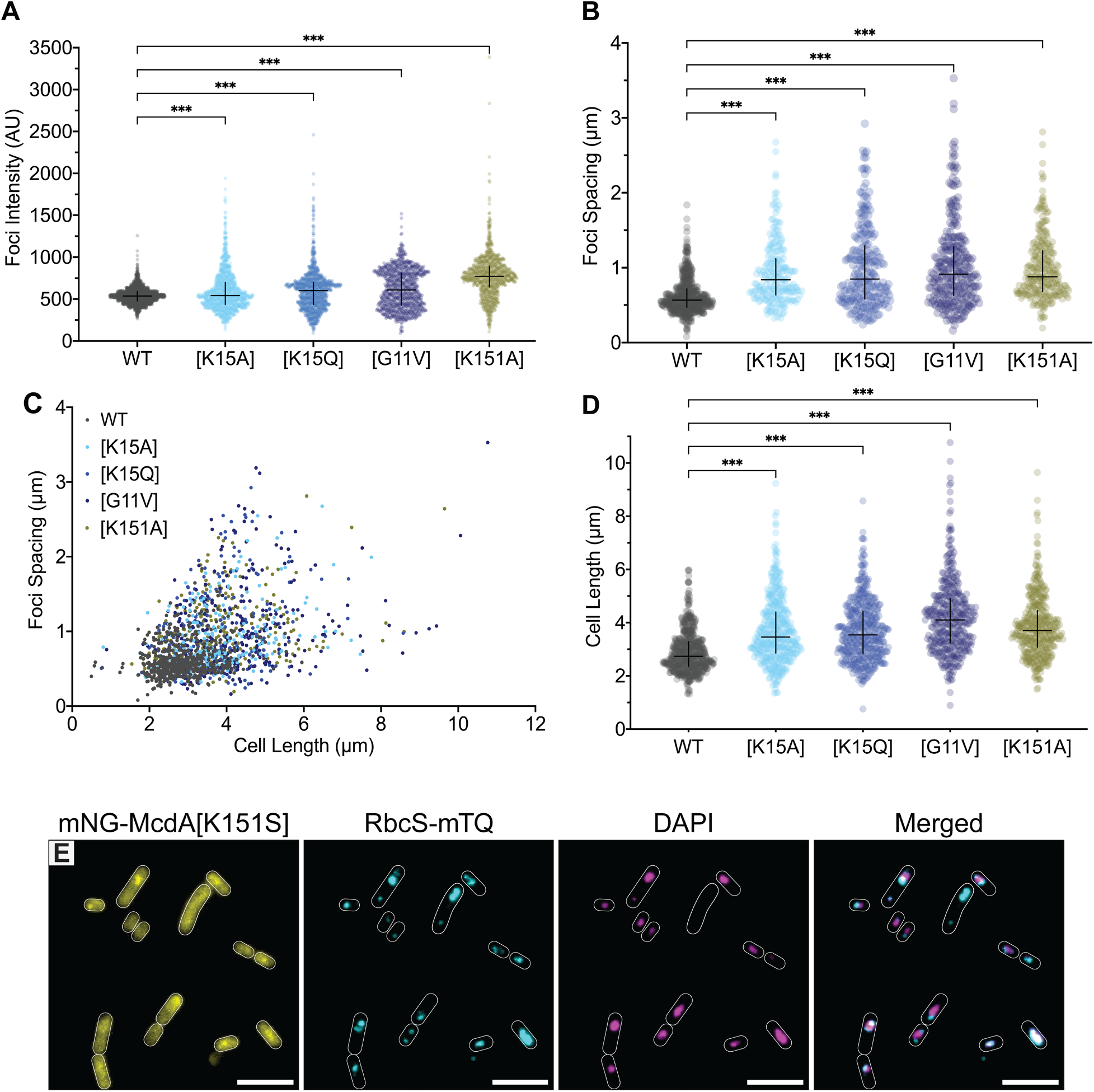
**(A)** Comparison of carboxysome foci intensity for specified strains. (Arbitrary Units = AU). WT *n* = 1925 foci; *n* > 1000 foci per mutant strain. **(B)** Comparison of carboxysome foci spacing of specified cell strains. **(C)** Distribution of carboxysome spacing as a function of cell length in the specified strains. For (**B**) and (**C**): WT *n* = 558 cells; n > 220 cells per mutant strain. **(D)** Comparison between the cell length of specified mNG-McdA mutants cell strains. WT *n* = 561 cells; *n* > 365 cells per mutant strain. (**E**) Microscopy images of mNG-McdA[K151S] (yellow), carboxysome foci (cyan) and DAPI-stained nucleoid (magenta) when treated with ciprofloxacin. Merged image shows carboxysome and DAPI signals.

## SUPPLEMENTARY MOVIE LEGEND

**Movie S1:** Live-cell fluorescence microscopy of wildtype *S. elongatus* cells (3 representative cells) treated with ciprofloxacin. mNG-McdA (yellow) continues to oscillate on ciprofloxacin-compacted nucleoids (DAPI) and carboxysomes (cyan) are still distributed across the compacted nucleoid. Movie was taken at 30 seconds per frame. Playback at 11 fps (330x real-time).

## SUPPLEMENTARY TABLES

**Table S1:**
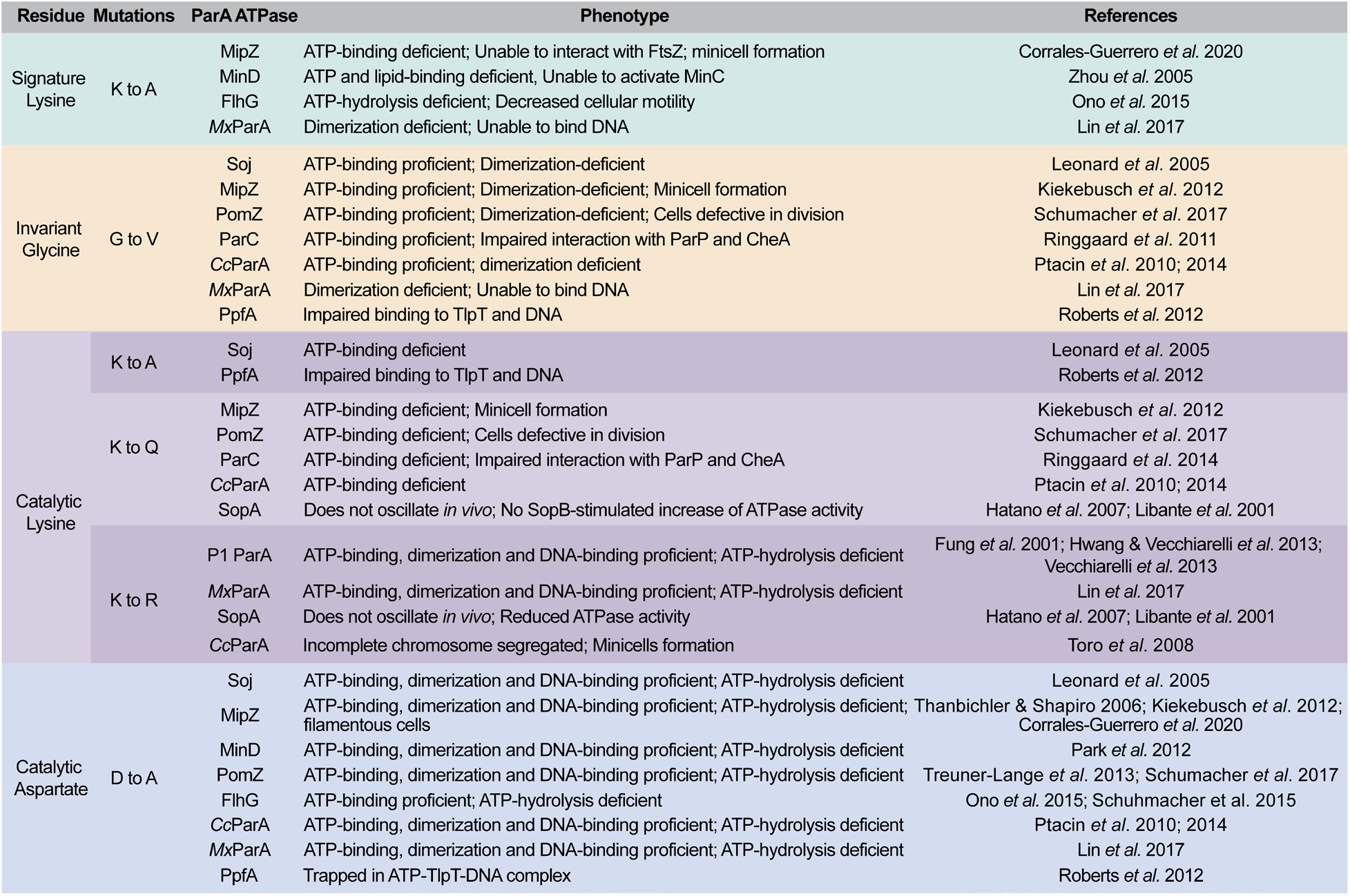
Detailed summary of ATP-binding pocket mutations studied in ParA family members and their associated phenotypes; Cc: *Caulobacter crescentus*, Mx: *Myxococcus xanthus.*

**Table S2:**
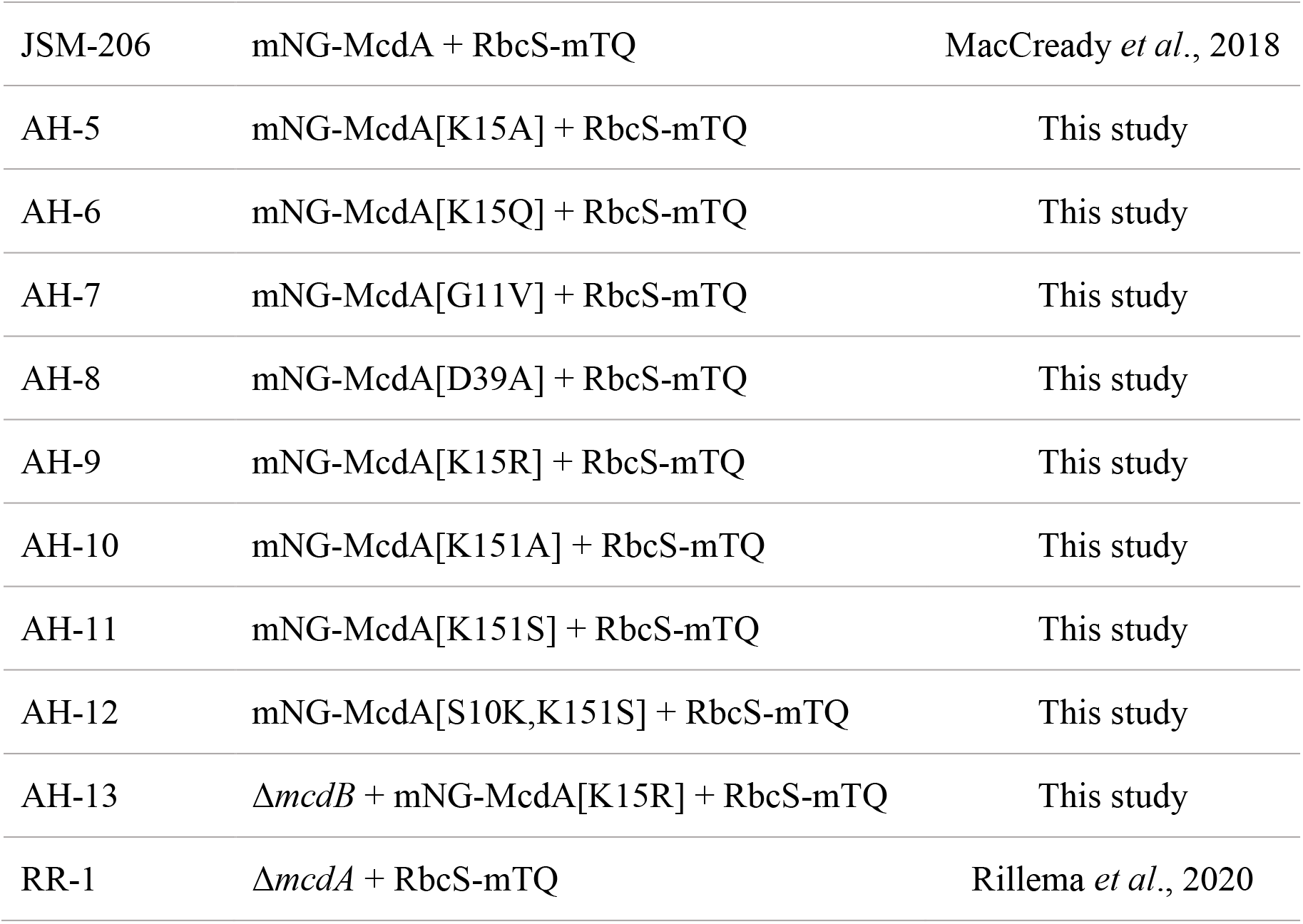
Cyanobacterial strains used in this study.

